# RADSuite: Improving Species-Level Identification in Microbiome Research

**DOI:** 10.64898/2026.11.18.767742

**Authors:** Rachel Christensen, Jakob Young, Myesha Gilliland, Remington Manning-Neal, Howard Fan, Sean Lu, Samuel Payne, W. Evan Johnson

## Abstract

In the last twenty years, the microbiome has become a central focus of biomedical research, with amplicon sequencing of the 16S rRNA gene considered the industry standard for taxon identification of prokaryotes due to its cost-effectiveness. However, common target regions in 16S amplicons often do not uniquely distinguish closely related species, restricting many analyses to the genus level. Additionally, curated databases such as NCBI’s 16S curated database often do not contain all intragenomic variants of the 16S gene and lack annotation of the 16S genomic composition and gene structure. To address these limitations, our team created RADSuite, a collection of tools which includes a more robust database of 16S gene copies (RADlib), an application that identifies which V regions will allow for species level identification (RADx), and a step-by-step guide for how to integrate RADlib within existing metagenomic profiling software tools. Here we demonstrate the utility of using RADSuite as a guide for designing 16S amplicon sequencing experiments. For example, RADx can be used to recommend 16S components that enable species-level resolution for most microbes of interest, even when industry-standard default amplicon choices (V3/V4) are insufficient. These findings suggest that targeted variable region selection is critical for reliable species identification. As researchers use RADSuite to improve the quality of their experiments, we anticipate improved experimental accuracy in fields of microbiome research from human health to environmental conservation.

## Introduction

Over the past two decades, interest in the microbiome has grown significantly, driven by the recognition that microbial communities shape both their surrounding environments and the health of their hosts. As a result, microbiome research now spans fields as diverse as pharmacology, ecology, nutrition, and pathogenesis [1–4]. A microbiome is the collection of all microorganisms living at a specific site: e.g. the soil at the base of a tree, a drop of ocean water, or locations on the human body. In fact, different parts of the human body, including the skin, intestines, mouth, etc., have distinct microbiomes [5]. Even regions previously thought to be sterile in healthy humans, such as the lungs, have been shown to have active microbiomes [6]. In studying human microbiomes, researchers have found that their compositions have a significant effect on many health concerns, including malnutrition, the progression of inflammatory diseases, and the metabolism of drugs [7–9]. In human microbiome research, as in all microbiome research, one of the primary goals is the identification of the individual species that exist in the microbiome. This is because even closely related species can exhibit distinct functional roles, leading to drastically different health outcomes [10–12]. The methods used to conduct microbiome research therefore prioritize improving researchers’ abilities to identify taxa at an individual level.

The current industry standard for taxonomic identification of prokaryotes in microbiome research is amplicon sequencing of the 16S rRNA gene [13,14]. The 16S rRNA gene is a component of the small subunit of the ribosome, and is about 1500 base pairs long. Because the ribosome is responsible for the translation of RNA into protein, the 16S gene is required for basic cellular function and is therefore present in all prokaryotes, making it an ideal target for species identification [15–17]. While the same is true for other essential genes, the structure of the 16S gene includes ten conserved regions and nine variable regions (V regions) which facilitate amplification based sequencing [14]. The conserved regions of 16S provide target sites for the primers used for amplification, while differences in the V regions allow for grouping and taxonomic assignment. When functioning optimally, this taxonomic assignment can be highly specific, allowing for species-level identification of microbes in a sample.

Despite the advantages of 16S amplicon sequencing, there are three primary issues which make it difficult for researchers to achieve species-level specificity, the first of which is limited data collection from the sample of interest. This is driven by cost-related factors; instead of sequencing the entire 16S gene, researchers often save costs by amplifying and sequencing only a few V regions. Partial sequencing is common because Illumina sequencing, the most widely used platform, is only capable of producing sequences <300 bases in length, less than the length of the 16S gene. The most commonly sequenced regions are either just the fourth V region or the third and fourth V regions together (denoted V3 and V4) [18,19]. This is because primers for the V3/V4 regions are the most readily available; Illumina even provides a specific 16S sequencing procedure for those regions alone [14]. Unfortunately, by only comparing select V regions, like V4 or V3/V4, instead of whole 16S rRNA genes, researchers are limited in their ability to discriminate between closely related organisms that share identical sequences in those regions. In fact, the conservation of the 16S V4 region is so extensive between related organisms that it is largely unhelpful in classification beyond the genus level [20]. Several methods have been developed to improve strain-level identification from 16S data [21]. Nevertheless, because closely related strains often have identical or nearly identical V region sequences, many of them still cannot be distinguished.

Even if researchers use optimal portions of the 16S gene, two issues remain; the content of and access to current 16S databases. First, curated databases, such as NCBI’s 16S rRNA curated database [22], Greengenes2’s 16S rRNA database [23], and SILVA’s rRNA database [24], do not include all paralogous copies of 16S rRNA for each species. A single microbial species possesses between one and fifteen copies of the 16S rRNA gene, with an average copy number of 4.2 [25]. Between copies in a single species, genetic variation tends to be small (mean intragenomic divergence 0.55%), although some outlier species can have as much as 5% intragenomic divergence [26,27]. When databases use only a few 16S copies per species, species which share identical sequences in those copies become indistinguishable from one another. Incorporating all copies of the gene in a reference database allows the variation between these copies to be exploited for more specific species identification [28]. Finally, while there are many databases with valuable information currently being maintained, it can be overwhelming for researchers to access and understand the information in these databases. This is exacerbated for researchers who are unfamiliar with coding tools and documentation.

To address the issues of species-level identification, database curation, and accessibility, our team created RADSuite. RADSuite is a collection of tools containing a more thorough database of 16S gene copies, an application that identifies the V regions which allow for species level identification, and a step-by-step guide for how to integrate our improved database with the existing analysis pipeline MetaScope. The database, called RADlib (Regional Alignment Database Library), includes all available copies of 16S genes from NCBI RefSeq’s prokaryotic whole genome database. The Rshiny application RAD Explorer (RADx) allows users to input organisms of interest and returns an interactive display of all sequences for each V region, along with information about which species can be uniquely identified when sequencing user-selected V regions. Users can then follow the step-by-step instructions included in the RADx interface to analyze sample data using the MetaScope pipeline with RADlib as the reference database. These tools will aid researchers in planning experiments which will result in species-level specificity, and thus more accurate results.

In this paper we discuss the databases and packages that comprise RADSuite. We describe the creation and contents of RADlib, the intended use cases for RADx, and available functionality of the RADalign package. We analyze the ability of RADSuite to provide a more thorough curated database, assist researchers in planning experiments, and improve database and pipeline accessibility for researchers at all levels of coding experience.

## Methods

### Structure Overview

RADSuite is a Github organization containing three main subsections, depicted in Figure 1 for ease of reference. RADlib, short for RADlibrary, comprises two databases: RADlib16S, which contains whole 16S rRNA genes, and RADlibVR, which contains the variable regions identified in RADlib16S. The data for RADlib is sourced from whole-genome shotgun sequences (wgs) stored in NCBI’s RefSeq database [29]. It also includes the Nextflow pipeline used to create RADlib and the documentation required to create an updated version of RADlib if the user desires. RADalign is an R package with two main functions. First, it extracts data from RADlib and performs analyses such as multiple sequence alignment (msa) and identification of identical sequence groups. Second, a section of the package titled RADintegrate extracts data from RADlib and downloads it in files compatible with the MetaScope analysis pipeline. Both use cases are made accessible through the final subsection, RADexplorer (RADx). RADx is an R Shiny application that displays the data and analyses created by RADalign for users to explore. Each of these subsections is described in greater detail below.

**Figure 1.**
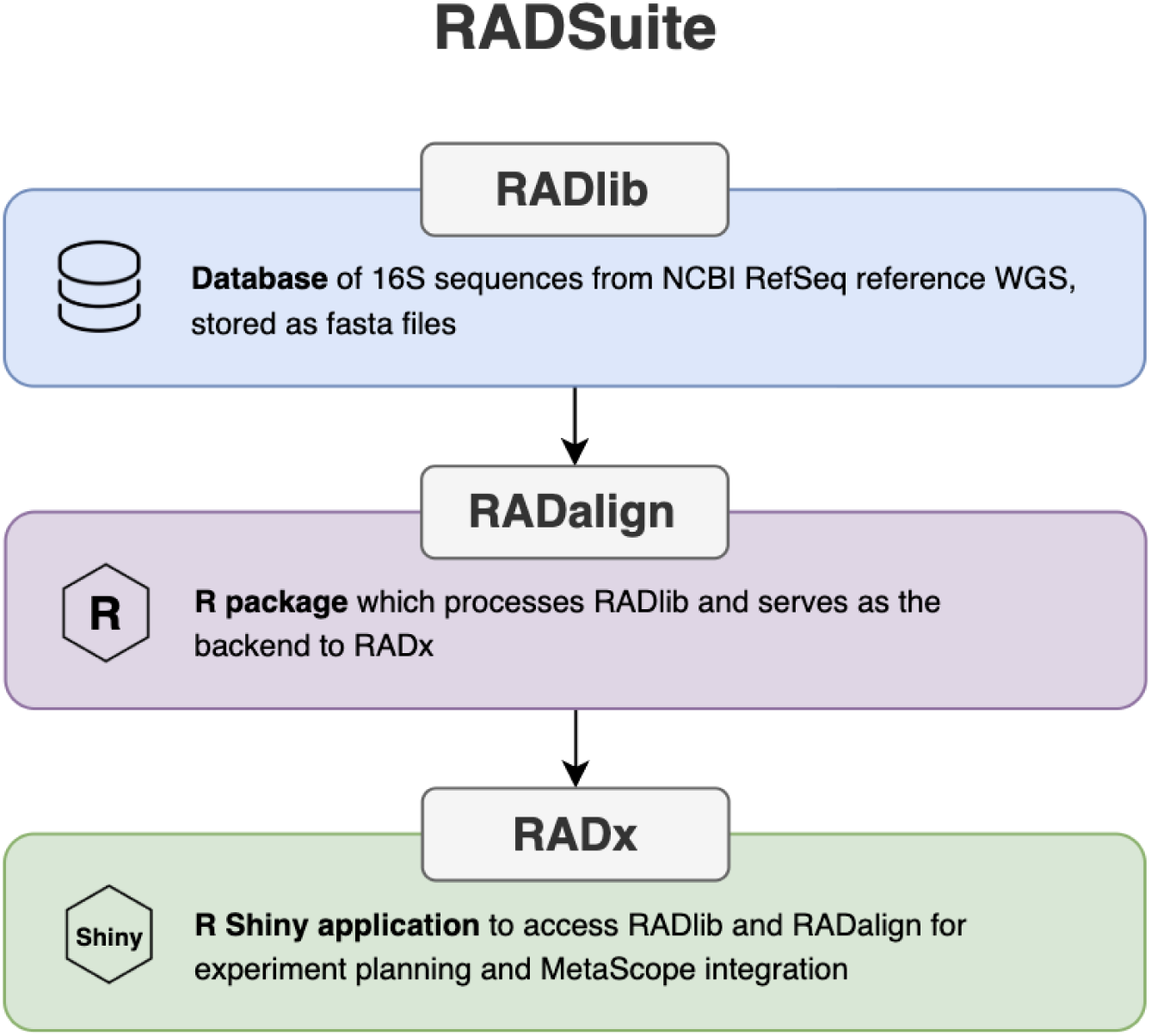
Basic structure of RADSuite. RADSuite consists of three main sections. RADlib is the database, created from whole-genome shotgun sequences (wgs) stored in NCBI’s RefSeq database. RADalign processes the data stored in RADlib, performs analyses, and outputs the data displayed in RADexplorer (RADx). RADx is an R Shiny application that displays the data and analyses created by the other two sections.

### RADlib16S and RADlibVR Curation and Analysis

To design a database that would allow for user exploration of copy number and variants within a species, we sourced our 16S rRNA genes from NCBI’s RefSeq wgs database [30]. All sequences are sourced from either the Bacteria or Archaea domains. A Nextflow pipeline [31] was designed to source reference genomes from RefSeq as well as their annotations created by their prokaryotic annotation pipeline [32] to streamline the 16S gene identification and extraction (Fig. S1). The annotated files were filtered for any lines annotating the 16S rRNA gene and passed, along with the wgs, to the Bedtools getFasta cli tool [33] to extract the genes. Each sequence header in the resulting FASTA file was then labeled with the locus tag, assigned by NCBI, the organism’s taxonomic id, and the organism’s name. Duplicate sequences were maintained for future use by RADx. The sequences were combined into a single FASTA file called RADlib16S.

In addition to our 16S rRNA database, we designed a database containing only the V regions within each 16S rRNA gene. To create this database we used a tool by Alessio Milanese to extract V regions [34]. The tool aligns each sequence against a reference 16S rRNA gene, *Escherichia coli K 12*. Alignment was performed using a pregenerated bacterial covariance model from the Rfam from June 2013 [35] using Infernal’s cmalign tool (v1.1.5) [36]. After alignment, the V regions of the reference sequence were identified and the region on the query sequence that aligned with the V region on the reference sequence was extracted and annotated as the corresponding V region. We then added headers of each V region sequence containing the same information as the 16S rRNA database but with the addition of a V region label. The resulting FASTA file was called RADlibVR.

### RADalign Design and Implementation

RADalign is an R package designed to use the RADlib reference library to explore the variation across 16S rRNA gene copies in microbial taxa by aligning 16S V regions. RADalign’s functionality begins with a list of user-selected taxa to explore. All V regions for the taxa are extracted from RADlibVR, then the sequences for each V region are aligned using the ClustalOmega algorithm contained in the msa Bioconductor package [37,38]. Sequences are then grouped according to which sequences shared identical alignments and assigned a V region sequence ID (seqID) unique to their group. The first output of RADalign is then produced: a csv or dataframe identifying each V region sequence by the seqID. Additional functions are provided that filter the output to only selected V regions, create a summary ID indicating which seqIDs are found in each species, or create a summary ID indicating which species are identifiable when sequencing selected V regions. Each of these functions produces an output file and can return a dataframe as requested by the user.

RADintegrate is a subset of functions and an instructions file included in the RADalign R package. These functions allow users to download the necessary files to use the MetaScope pipeline to analyze sample data. Inputs to these functions are passed in through the RADx interface and include download of the full or partial RADlib, which species to include in a partial RADlib, and the download location, as well as an option to use an updated version of RADlib (see S1) or a fasta file which meets the RADlib formatting specifications. Simple instructions, written for those with little to no code experience, describe how to integrate the downloaded reference database with MetaScope. These instructions are included in the supplementary folder of RADalign, and are visible in the RADx display. A tutorial run of MetaScope with an example dataset is included in the instructions.

The primary function of RADalign is to provide a backend for the RADx app, with secondary functionality allowing users who are comfortable with coding in R to access the data passed to RADx. The summary IDs and filtered outputs correlate with display options in RADx and are most easily interpreted using that format. For users who want to access the data directly using R code, additional information on package functionality can be found using roxygen documentation by running the R command “?function” or “help(function)”. All code and documentation can be found in the RADalign repository found in the RADSuite github linked below.

### RADx Design and Implementation

RADx was built as a R Shiny application that makes RADlib data, RADalign functionality, and MetaScope integration accessible to biologists across all levels of programming experience (see Figure 2) [39]. Users select taxa of interest and one or more V regions, which are passed internally to RADlib and RADalign to retrieve the relevant sequences and perform groupings analyses. Outputs from each RAD package are rendered as an interpretable tile plot, built in ggplot and converted into an interactive figure using plotly [40,41]. A summary view was implemented that collapses each taxon’s gene copies into a single signature per V region, allowing for quick interspecies comparison of V regions. In addition, a detailed view was implemented that displays the V regions in all individual gene copies for each taxon, ideal for intraspecies gene copy analysis. Visual elements, such as green checkmarks and red brackets, were used to display the results of RADalign grouping analyses and communicate to the user the efficacy of the selected V regions. On the RADx explore page, additional functionality like “Locate Taxa” were added to allow the user to browse and filter the figure for the main taxa they are interested in. An adaptive instructions screen on RADx was built so that RADlib data can be downloaded and integrated with MetaScope by users.

**Figure 2.**
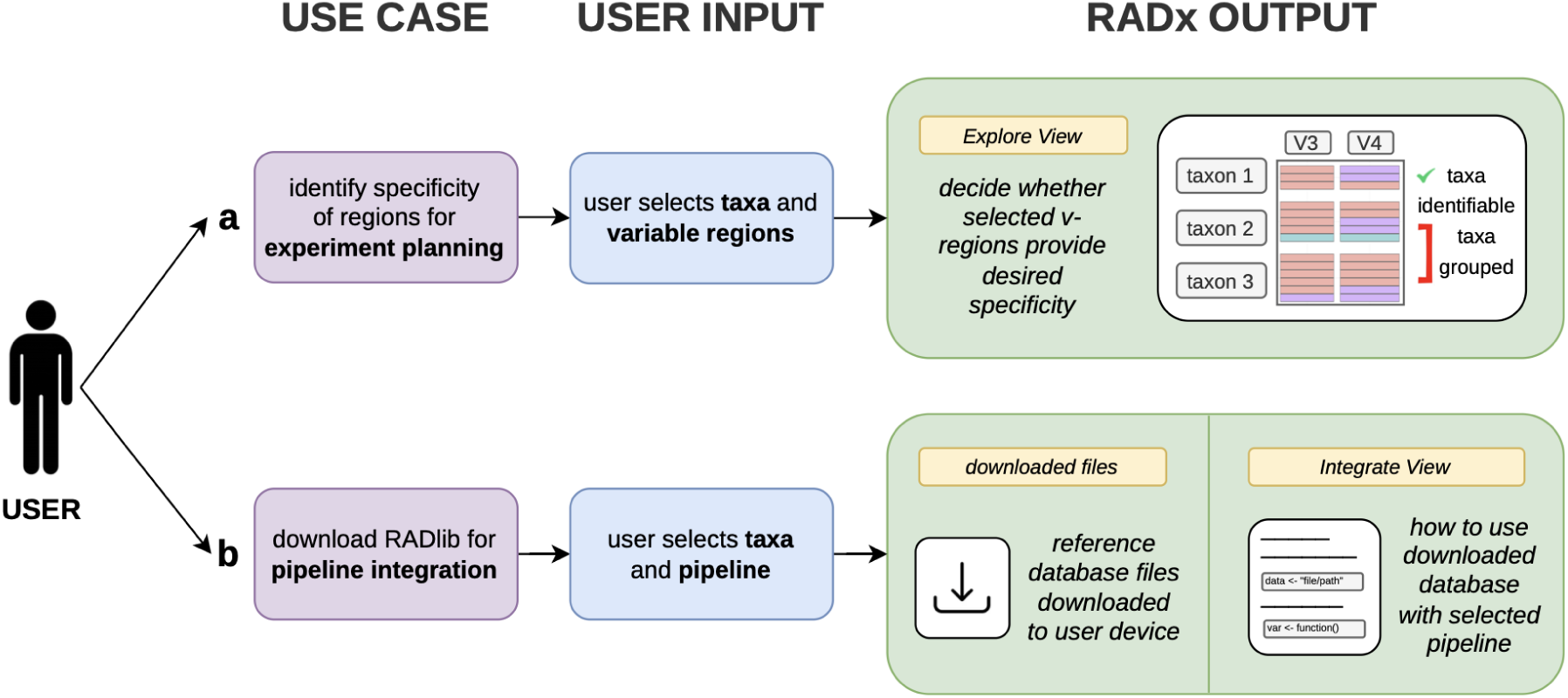
Use cases for RADexplorer. (**a**) To compare taxa of interest, the user selects the taxa from a list of all genera and species included in the RADlib sequence library. The RADalign package filters the selected taxa out of RADlib, performs a multiple sequence alignment on each variable region (V region), and returns the results of the alignment to the RADexplorer app (RADx). In RADx’s all regions view, each gene copy present in a taxon is displayed as a colored bar. Bars with the same color indicate sequences that align exactly within each V region. Once selected taxa have been displayed with all V regions, the user can select specific V regions to filter the display. RADx will also display which taxa are uniquely identifiable with the selected V regions and which taxa will be grouped due to sharing V region sequences, indicated by green check marks and red brackets respectively. (**b**) The user can also use RADx to integrate RADlib with the MetaScope analysis pipeline. RADalign downloads either the full RADlib or a user-selected set of sequences from RADlib, and displays instructions for how to use the downloaded files as the reference database for the MetaScope pipeline.

### Accessibility

All code for RADx, RADalign, RADlib creation, and figure regeneration (including access to data sets used) can be found on Github: https://github.com/RADSuite. The authors request that any use of the provided resources cite this manuscript.

## Results

To address the challenges in 16S amplicon research, we created RADSuite, a Github organization that grants users access to our curated database, facilitates improved experiment planning, and provides instructions for integrating the database with the MetaScope analysis pipeline. Our curated database, RADlib, addresses the problem of data missing from current databases by including more 16S references per species and by annotating the V regions within each 16S reference. The RADalign and RADx packages work together to alleviate the difficulty of identifying taxa at a species level by providing a tool for experiment planning that allows users to view which V regions can be used to differentiate species within their selected species group. The RADalign and RADx packages also improve data accessibility by providing clear instructions designed for users with little coding experience to be able to use the MetaScope analysis pipeline. User access of these two RADx features are outlined in Figure 2. Our improved database, experiment planning, and pipeline integration are discussed in the following sections.

### Improved Reference Counts and Annotation in RADlib

RADlib16S on average has more 16S rRNA sequences per species as compared to NCBI’s 16S reference database. This is because RADlib16S maintains all available paralogous genes or copies of 16S rRNA sequences found within an organism, including identical copies. The comparison between copy numbers in the databases for five frequently-studied microbiome species can be seen in Figure 3. As an example, *Escherichia coli* (*E. coli*) genomes contain on average 7 copies of the 16S rRNA gene with some of those genes having distinct variable regions. The NCBI 16S database contains 8 *E. coli* reference genomes; for 7 of those genomes the database contains only 1 16S gene. This accounts for the 14 total 16S genes for *E. coli* in our database. Even though 7 of the references were not fully annotated, RADlib still has significantly more total references than NCBI’s curated database which has only 3 references, each of which are distinct. One exception to RADlib having more references for a given taxa is *Enterococcus faecalis* (*E. faecalis*). In this species, RADlib has one fewer reference than NCBI’s curated database. The reason for this discrepancy is that NCBI’s curated database occasionally draws from some non-reference organisms, resulting in their higher number of references in this case.

**Figure 3.**
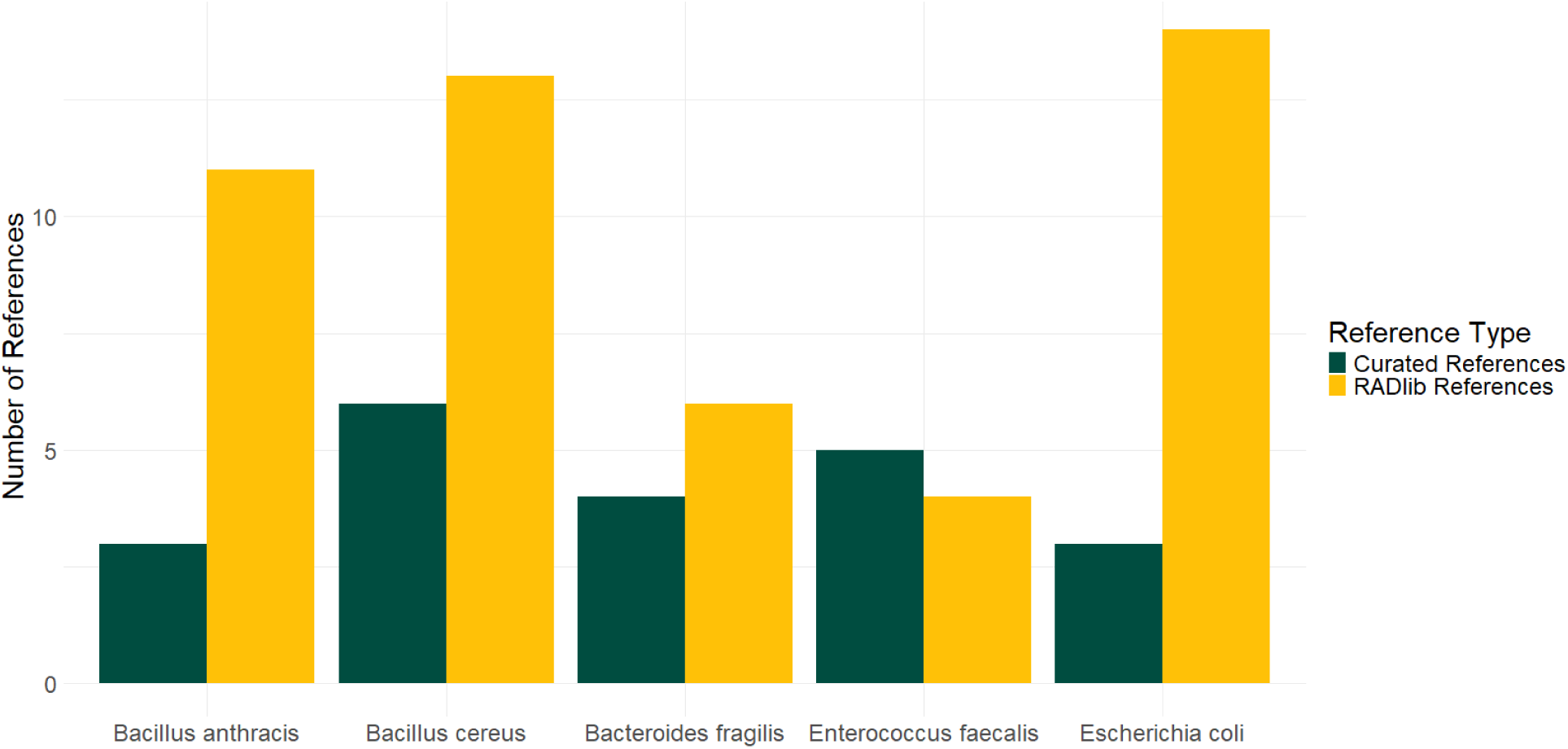
Number of 16S rRNA gene sequences in NCBI’s 16S curated database and in RADlib for a given species. Because RADlib draws from all whole genomes available in NCBI RefSeq’s whole genome database, it contains more 16S sequences per species on average than NCBI’s 16S reference database. NCBI’s reference database does not contain duplicate sequences from within the same species while RADlib does. However, NCBI draws from some non-reference organisms for their 16S database, accounting for the greater number of *Enterococcus faecalis* references in NCBI’s 16S database.

There are notable limitations to RADlib’s method of sourcing 16S rRNA genes. First, drawing strictly from the NCBI wgs database reference genomes does not capture the diversity within some subspecies and strains due to the limitations of what NCBI’s wgs database contains. Second, NCBI’s wgs database does not contain fully reconstructed genomes for each organism, such as some *E. coli* strains. Other available databases may contain more 16S rRNA genes than RADlib16S because they draw from multiple sources [42]. However, what makes RADlib unique is RADlibVR. Many 16S rRNA databases do not contain annotations of the V regions contained within their database, but RADlibVR provides this additional information. This allows researchers to better plan their experiments by identifying which V regions they should select to provide the highest taxonomic resolution possible using RADlib16S. This provides researchers an opportunity to determine which primers are needed to perform their amplification, as well as address any potential financial, sequence error rate, or other limitations that may come from their desired primer combinations.

### Designing Effective Amplicon Experiments with RADx

RADx produces interpretable visualizations of 16S rRNA gene copy variation between taxa. Differences in V region sequences that would otherwise require direct sequence comparison are made visible on the Explore page. The interface displays RADalign grouping analysis, making it straightforward to identify which V regions resolve taxa of interest down to the species level. This is particularly evident between closely related taxa, where the choice of V region has substantial consequences on taxonomic resolution.

As shown in Figure 4, common amplicon regions fail to achieve species-level resolution in the Escherichia species group. RADx makes this limitation visible and avoidable. Specifically, V3 fails to differentiate any species in the species group. V4 performs marginally better, resolving two species while leaving the remainder indistinguishable from one another. In contrast, V1 by itself can identify every *Escherichia* species included in RADlib down to the species level. This result demonstrates that V3 and V4, despite their widespread use in amplicon studies, are not always the best choice for resolution. For some taxa, they can be among the least effective regions available.

**Figure 4.**
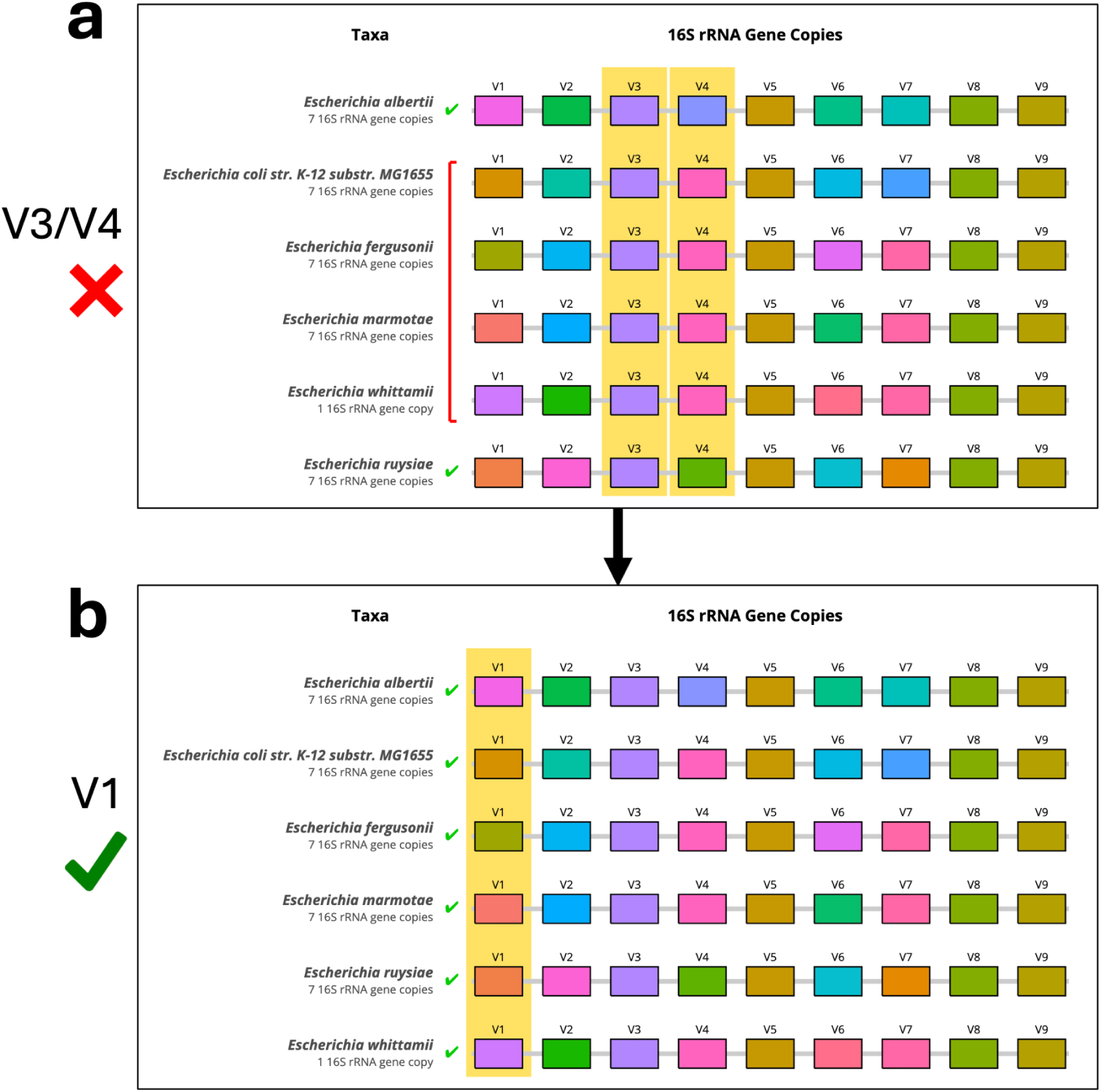
Screenshots of RADx distinguishing Escherichia species with V3/V4 (a) versus V1 (b). When the V3 and V4 amplicons are selected together, only two *Escherichia* species are resolved: *Escherichia albertii* and *Escherichia ruysiae*. When V1 is selected by itself, all *Escherichia* species are identified to the species level.

The limitations of the V3/V4 regions are not unique to *Escherichia*; RADx-identified amplicon combinations consistently outperform V4 and V3/V4 (Fig. 5). Across the six tested genera, V4 distinguished only some of the available species in each genus, and V3/V4 provided minimal improvement. In contrast, RADx-recommended V region combinations could identify most if not all available species in each genus. Importantly, the best combination of amplicon regions varied by genus. This demonstrates that no single V region combination is universally optimal. Defaulting to a standard region without prior evaluation may leave an experiment incapable of answering the biological question it was designed to address. Instead of requiring a dedicated study on a genus to ensure species-level resolution via amplicon sequencing, RADx-recommended combinations give researchers confidence that their taxa of interest will be identified.

**Figure 5.**
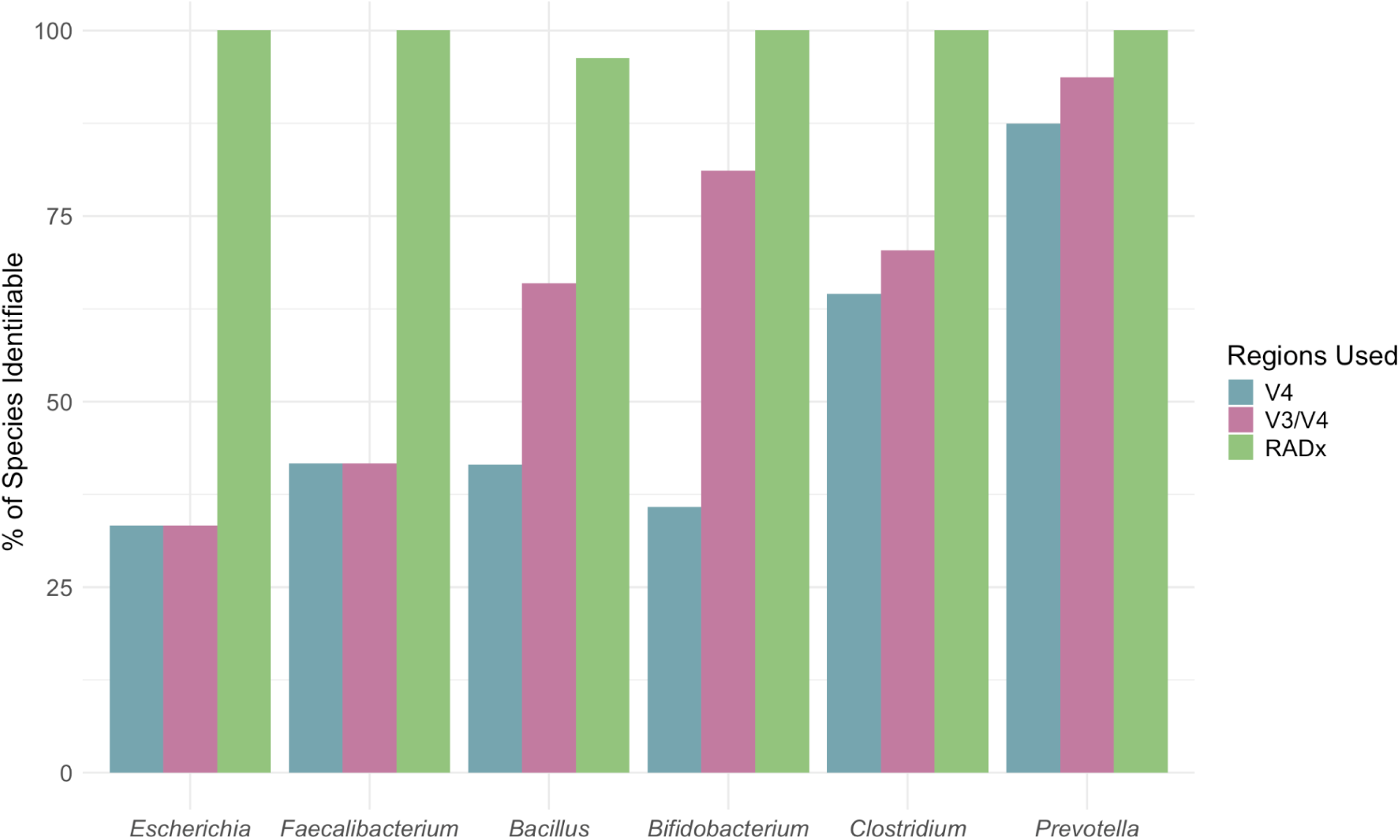
Comparison of species-level resolution achieved by V4, V3/V4, and RADx-recommended amplicon regions across six genera. Each genus is listed with three bars representing the percentage of species in the genus identifiable with V4 alone, combined V3/V4, and the regions recommended by RADx. The regions recommended by RADx for each genera are: V1 (*Escherichia*), V1/V6 (*Faecalibacterium*), V1-V4/V8-V9 (*Bacillus*), V6-V9 (*Bifidobacterium*), V1-V4 (*Clostridium*), and V1/V9 (*Prevotella*).

In addition, RADx identifies cases where species-level resolution using 16S amplicon sequencing is impossible. For example, three *Bacillus* species – *B. albus, B. basilensis,* and *B. dicomae* – have completely identical V regions across all 16S gene copies. In this case, no combination of V regions can distinguish one species from another. Even sequencing the entire 16S gene would fall short of discriminating between these three closely related species. RADx makes this limitation visible, demonstrating that researchers can avoid spending time and resources on sequencing methods that will not be able to identify species of interest.

### Pipeline Integration: Sample Analysis using RADlib

We created RADintegrate to ensure that RADlib would not only enable scientists to determine the ideal combination of 16S V regions to sequence, but would also be accessible to researchers as a reference database for analysis of sample data. Using RADlib as the reference database requires some adjustments to the usual MetaScope protocol, which could prove difficult for those unfamiliar with code and documentation. To address this concern and widen the accessibility of these tools, we created instructions which are designed to be comprehensible even to those with little coding experience. These simple, easy-to-follow, copy-and-paste instructions teach researchers how to run the powerful analysis of the MetaScope pipeline on their own data, using RADlib. Researchers wishing to run the MetaScope pipeline on their own device can access the RADintegrate instructions on the RADx Integrate page. For analysis of larger sample datasets which require the use of a supercomputer, adjusted instructions which do not rely on the RADx interface are provided as a markdown file in the supplementary folder of the RADalign package.

## Discussion

With RADSuite, researchers with any level of programming experience can now access, interpret, and act according to 16S gene copy data. While the underlying data has existed for years, without a way to understand and interact with it many researchers lacked a practical way to apply it to their work. In microbiome research, where ineffective V region selection can leave many species undetected and change the conclusions drawn from a dataset, having this data visible and accessible is a necessity. Now, with RADx, what once would have required hours of sequence analysis for each experiment is now displayed in a single interactive view that any researcher can interpret. Because of that visualization, every researcher has the ability to make an informed V region selection and understand the genetic basis of that decision.

In addition, modern classification tools are designed to use databases that contain only distinct sequences within an organism’s genetic material. To allow these databases to be distributed to researchers, removing redundant sequences allows the databases to remain small and have more diversity. This design philosophy obfuscates the reality that prokaryotes can have more than one identical copy of the 16S gene in their genetic material. Our hope is that RADlib16S brings to light a need for databases that contain information about copy numbers so future classification algorithms can better discriminate what organisms are in a sample and in what proportion. Future research is still needed to determine the optimal database format and algorithmic approach to accomplish this goal.

RADx also has the potential to increase the cost efficiency of species-level identification in microbiome research across many fields of study. Partial sequencing of the 16S gene has been generally preferred to whole genome sequencing due to cost and time considerations, but at a significant cost to the ability of researchers to classify sequences with species-level specificity. This limitation has direct clinical consequences. For example, specific microbial species such as *Enterococcus faecalis* and *Eggerthella lenta* have been shown to reduce the bioavailability of a key Parkinson’s disease medication, a finding that would be impossible to act on without reliable species-level identification [9]. By using RADx to plan experiments, researchers can select the optimal V regions to amplify, thereby improving species identification in their target taxa while preserving the cost-efficiency of only amplifying part of the 16S gene. We anticipate that RADx will make species-level identification possible for many more researchers. This improvement will in turn have a positive impact on research in the many fields impacted by the study of the microbiome.

## Supplementary

**Figure S1.**
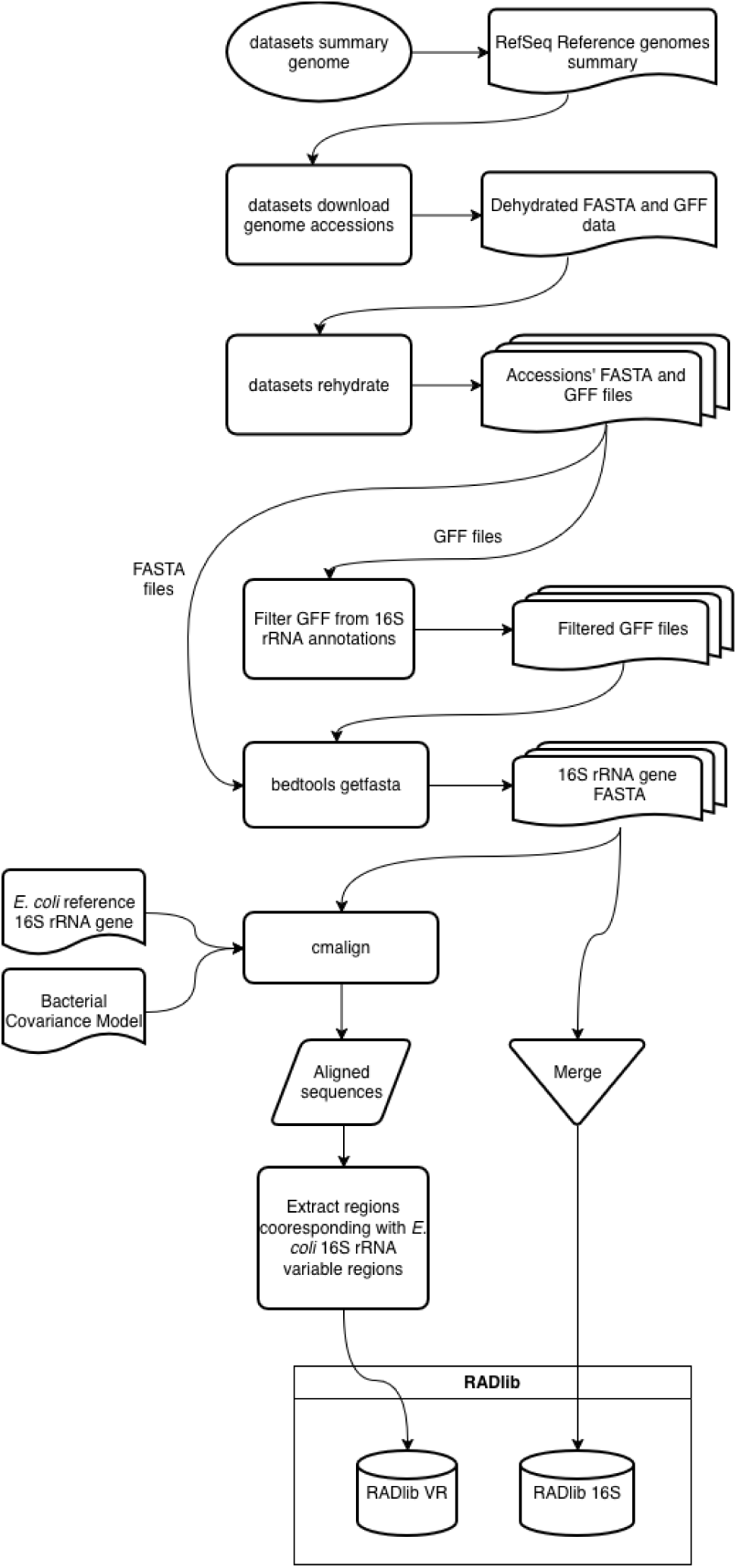
A workflow diagram of RADlib creation. Data is sourced from NCBI’s whole genome shotgun database and 16S sequences are extracted. Cmalign and region extraction is performed by extract_regions_16S.

## Notes

https://github.com/RADSuite

### Competing Interest Statement

The authors have declared no competing interest.

